# Allosteric activation dictates PRC2 activity independent of its recruitment to chromatin

**DOI:** 10.1101/206383

**Authors:** Chul-Hwan Lee, Jia-Ray Yu, Sunil Kumar, Ying Jin, Syuzo Kaneko, Andrew D. Hamilton, Danny Reinberg

## Abstract

PRC2 is a therapeutic target for several types of cancers currently undergoing clinical trials. Its activity is regulated by a positive feedback loop whereby its terminal enzymatic product, H3K27me3, is specifically recognized and bound by an aromatic cage present in its EED subunit. The ensuing allosteric activation of the complex stimulates H3K27me3 deposition on chromatin. Here, we report a step-wise feedback mechanism entailing key residues within distinctive interfacing motifs of EZH2 or EED that are found mutated in cancers and/or Weaver syndrome. PRC2 harboring these EZH2 or EED mutants manifest little activity *in vivo* but, unexpectedly, exhibited similar chromatin association as wild-type PRC2, indicating an uncoupling of PRC2 activity and recruitment. With genetic and chemical tools, we further demonstrated that targeting allosteric activation overrode the gain-of-function effect of EZH2^Y646X^ oncogenic mutations. These results revealed critical implications to the regulation and biology of PRC2 and a novel vulnerability in tackling PRC2-addicted cancers.

## Introduction

Polycomb group (PcG) proteins are key epigenetic regulators that maintain transcriptional repression once established. During normal development, polycomb-mediated repression of lineage-specific genes contributes to the stability of cell identity and animal body plan formation^1–3^. Dysregulation of PcG proteins induces aberrant transcriptional programs that contribute to developmental diseases as well as cancer progression^4–7^. A case in point is Polycomb Repressive Complex 2 (PRC2). A hallmark of PRC2-mediated gene silencing is the methylation of histone H3 lysine 27 (H3K27), with H3K27me2/3 being associated with repressive chromatin^8^. The core complex of PRC2 comprises three subunits: EZH1/2 (Enhancer of Zeste Homolog 1 or 2), EED (embryonic ectoderm development), and SUZ12 (Suppressor of Zeste 12), that associates with the histone binding protein RbAp46/48 (Retinoblastoma-associated proteins 46 or 48)^9–15^. EZH1 and EZH2 are interchangeable subunits of PRC2 that harbor the histone lysine N-methyltransferase (HMT) activity through their respective SET domains, with the level of HMT activity exhibited by PRC2 being markedly elevated when comprising EZH2, relative to EZH1^16, and Lee et al, in submission^. The EZH1/2 catalytic SET domain exhibits an autoinhibitory state^17,18^, which is relieved upon EZH1/2 association with EED and SUZ12, prerequisite for HMT activity. In addition, PRC2 exists *in vivo* as a holo-complex, associating with different accessory proteins that regulate its activity and recruitment to chromatin^19–29^. One such protein is Jarid2 that assists in the recruitment of PRC2 to developmental loci in mouse embryonic stem cells (mESC) and, more importantly, enhances its HMT enzymatic activity^22,23,29–32^. In addition, a recent study has demonstrated that Polycomb-like proteins (PCLs), particularly MTF2/PCL2, facilitate PRC2 recruitment to CpG islands in mESC^33^. Our previous work demonstrated that EED binds to H3K27me3 through its aromatic cage, leading to an allosteric activation of the complex, further assisting PRC2 in establishing repressive chromatin domains^34^. Of note, PRC2 association with Jarid2 results in Jarid2 being tri-methylated at lysine residue 116 (Jarid2-K116me3), which also stimulates PRC2 in a manner similar to that of H3K27me3, and this function is likely required for the *de novo* deposition of H3K27me3^30^.

In human cancers, PRC2 plays a pleiotropic and highly context-dependent role. Loss-of-function mutations and deletions of EZH2 or SUZ12 have been found in multiple subtypes of T cell acute lymphoblastic leukemia (T-ALLs)^35,36^, and those of EED or SUZ12 were found in malignant peripheral nerve sheath tumors (MPNSTs)^35,37^. In addition, a vast reduction in H3K27me3 was observed in diffuse intrinsic pediatric glioma (DIPG) that harbors a K to M substitution at position 27 within any of the isoforms of histone H3 (H3K27M)^38,39^. On the other hand, EZH2 was found to be over-expressed in many other types of human cancers including breast, prostate, gastric, uterine, non-small cell lung cancers, and cutaneous melanoma^5,40^. Its high expression in these cancers was associated with poor prognosis^4,5,41–44^. Also, the “hotspot” gain-of-function mutations at residue Y646 within the catalytic SET domain of EZH2 were found in ~20% of diffuse large B-cell lymphoma (DLBCL), ~12% of follicular lymphoma, and ~1% of melanoma^35,45–48^. The Y646X (X=S,N,F,C, or H) mutations within the SET domain of EZH2 specifically switches the substrate preference of EZH2, thereby facilitating the conversion of H3K27 di-methylation to the tri-methylation state in peptide-based HMT assays *in vitro*^47,49^. Thus, the Y646X mutations of EZH2, are considered to be gain-of-function for H3K27me3 and EZH2^Y646X^ mutant cancers often manifest a dependency on PRC2/EZH2^50,51^. Therefore, EZH2 has been implicated as a therapeutic target in several types of cancer, including non-Hodgkin Lymphoma and DLBCL that rely on the activity of oncogenic EZH2^51,52^. Thus, EZH2 inhibitors were developed as competitive inhibitors of S-adenosylmethionine (SAM), the methyl-donor essential for the HMT activity, and several of these inhibitors are currently in clinical trials^51,53^. However, the therapeutic index of these SAM-competitive inhibitors is restricted by their imperfect pharmacological properties, including their short half-life, moderate to high clearance rate, and low permeability in cell-based assays^51,53,54^. Of note, in cellular models of DLBCL, *de novo* mutations at residues I109, Y111, and Y661 of EZH2 were found to prevent inhibitor binding and led to an acquired resistance to these inhibitors^55,56^. Therefore, the requirement for new types of EZH2 inhibitors is foreseeable. For example, new inhibitors targeting the cage of EED have been developed to abrogate the EED-H3K27me3 interaction and thereby prevent the initiation of H3K27me3-potentiated positive feedback activation of PRC2^57–61^.

Recently, the structures of PRC2 from human and a thermophile fungus, *Chaetomium thermophilum*, were solved^17,18^. The structure of the fungal PRC2 was shown in its basal as well as allosterically stimulated states, and that of human PRC2 in a stimulated state with a Jarid2-K116me3 peptide^17,18^. Additionally, the structure of a chimeric PRC2 consisting of EED and SUZ12 from human together with EZH2 from American chameleon (*Anolis carolinensis*) was solved in its basal state^62^. The structures of human and fungal PRC2 illustrated a major reorganization of the complex upon EED binding to H3K27me3 or Jarid2-K116me3, namely the stabilization of the SET-I domain (an α-helix of the catalytic SET domain) through its interaction with an α-helix of the Stimulatory Responsive Motif (SRM) of EZH2^16^.

Although previous structural studies have implicated the involvement of the SRM and SET-I domains of EZH2 in PRC2 allosteric activation, the structures of PRC2 were derived in complex with either the H3K27M oncohistone or a SAM competitive inhibitor which might conceal the dynamics of the SET domain during its allosteric activation^17,18,62^. Yet, whether PRC2 acts in the same fashion while in its native state *in vivo* is unclear. To better understand the molecular mechanisms underlying the allosteric activation of PRC2, we revisited the structural data of PRC2 and examined the genetic mutations of PRC2 found in human diseases. We further analyzed the key regions that undergo conformational remodeling during allosteric activation of PRC2 and pinpointed the functional significance of key residues by introducing disease-associated point mutations. The biological impact of the allosteric activation of PRC2 was then investigated in a variety of cellular models, including mESC, melanoma and DLBCL cells.

## Results

### A stepwise model for allosteric activation of PRC2

Previous studies demonstrated that EED, through its aromatic cage, binds to H3K27me3 or Jarid2-K116me3, leading to the allosteric activation of PRC2^30,34^. This enhanced activity entails a remodeling of the PRC2 subunits that orders the SRM domain of EZH2, based on recent structural data of PRC2^18^. Together, the structural and previous results favor a stepwise model for the allosteric activation of PRC2^18,30,34^, which involves an initial recognition (EED aromatic cage binding to H3K27me3/Jarid2-K116me3), an intermediate state (ordering of EZH2-SRM followed by the stabilization of the EZH2-SET-I domain) and a final step for substrate recognition and catalysis by a fully empowered SET domain (Figure 1A). Hereafter, we refer to the EED-H3K27me3 binding step as “recognition” and the deposition of PRC2 onto chromatin as “recruitment”.

**Figure 1.**
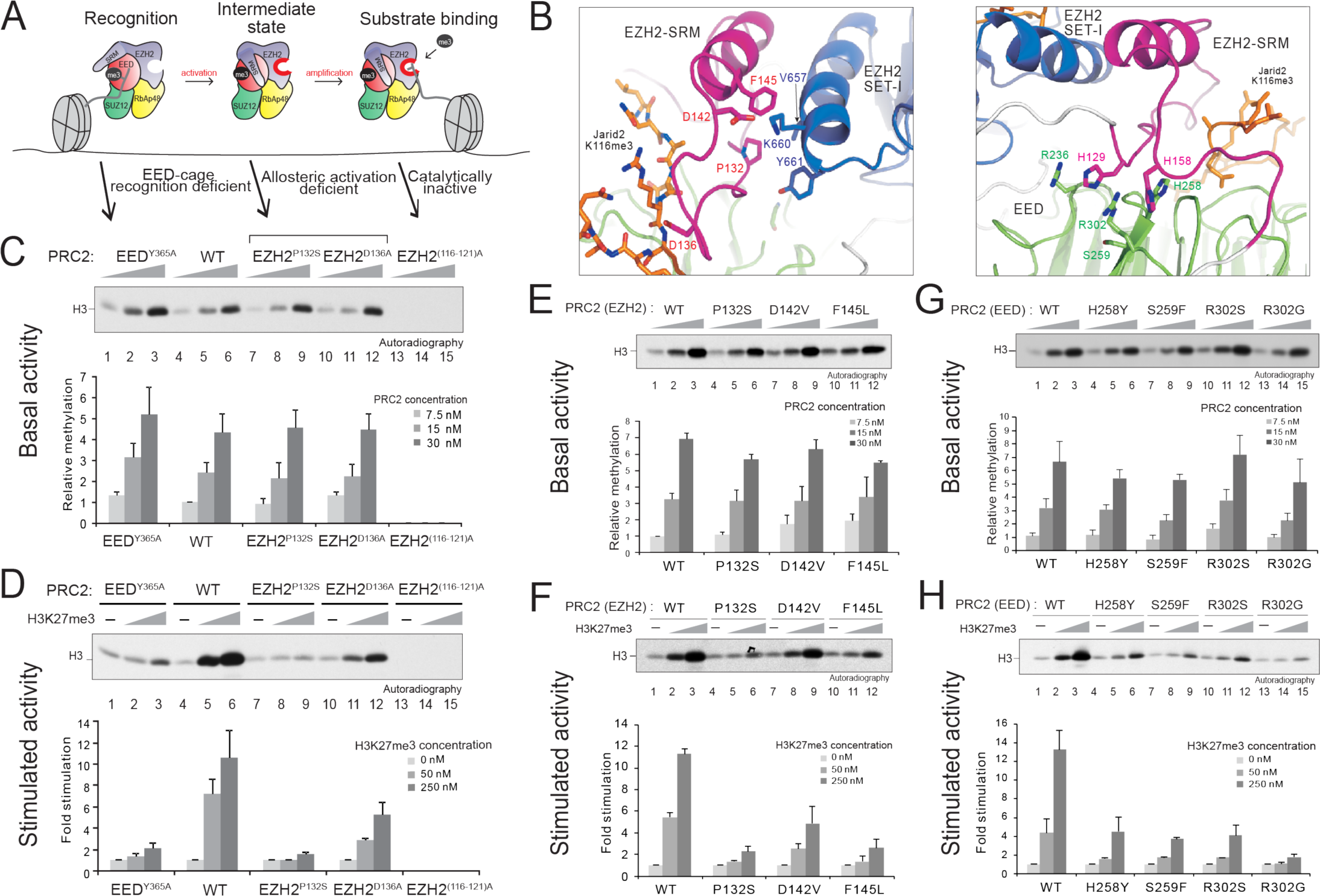
Identification of key EZH2 and EED residues required for allosteric activation of PRC2. (A) Illustration of the stepwise model of H3K27me3-potentiated positive feedback activation of PRC2, including the initial recognition step (EED binding to H3K27me3), the intermediate state (EZH2-SRM and SET-I ordering), and the substrate binding step (catalysis). Arrows indicate the corresponding PRC2 mutants for the dissection of each step within the model. (B) Structural depictions of the EZH2-SRM/SET-I (left) and the EZH2-SRM/EED interfaces (right) in the presence of the Jarid2-K116me3 peptide (amino acids 111-121), marked in orange. Residues that mediate interactions between the interfaces are highlighted. (modified from PDB:5HYN) (C-H) Histone methyltransferase assays. Details of the HMT assay conditions as well as the generation of the oligonucleosomes are described in Materials & Methods. (C-D) HMT assays using wild-type (WT) PRC2 or PRC2 containing an EED cage recognition mutant (EED^Y365A^), EZH2-SRM allosteric activation-deficient mutants (EZH2^P132S^ and EZH2^D136A^), or a catalytically inactive mutant (EZH2^(116-121)A^) with unmodified oligonucleosomes (150 nM) as substrate. (C) *Top*, a representative HMT assay measuring the basal activity of PRC2. HMT assays were performed with an increasing amount of PRC2 (7.5, 15, or 30 nM) in the absence of the H3K27me3 stimulatory peptide (amino acids 21-44) as illustrated in the panel. The levels of methylation on histone H3K27 are shown by autoradiography. (D) *Top*, representative assay measuring the stimulated activity of PRC2. HMT assays were performed using 15 nM PRC2 without or with an increasing amount of the H3K27me3 peptide (50, or 250 nM). (C-D) *Bottom*, Quantification of the relative amounts of ^3^H-SAM incorporated into histone H3 after 60 min incubation of the assays described in panel (C) (n=3 for each data point). (E-F) HMT assays using WT PRC2 or PRC2 containing EZH2-SRM mutants (EZH2^P132S^, EZH2^D142V^ or EZH2^F145L^). *Top*, Representative HMT assay measuring the basal activity (panel E) and the stimulated activity (panel F) of PRC2 with the experimental conditions described in panel (C-D). *Bottom*, Quantification of the relative amounts of ^3^H-SAM incorporated into histone H3 for the assays described in panel (E) (n=3 for each data point). (G-H) HMT assay using WT PRC2 or PRC2 containing EED anchoring mutants (EED^H258Y^, EED^S259F^, EED^R302S^, or EED^R302G^). *Top*, Representative HMT assay measuring the basal activity (panel G) and the stimulated activity (panel H) of PRC2 under the experimental conditions described in panel (C-D). *Bottom*, Quantification of the relative amounts of ^3^H-SAM incorporated into histone H3 for the assays described in panel (G) (n=3 for each data point).

### Specific mutations at the interface of EZH2-SRM/SET-I and that of EZH2-SRM/EED abrogate allosteric activation of PRC2

To test the model proposed above, we first scrutinized the structural features of the SRM domain of human EZH2. Upon EED binding to the Jarid2-K116me3 peptide, the EZH2-SRM formed a loop-α-helix-loop structure, with one side of the α-helix interacting with the SET-I domain of EZH2 and the other with Jarid2-K116me3 and EED (Figure 1B, *left*)^18^. Several residues of the α-helices within the SRM and SET-I formed hydrophobic interactions, including P132 and F145 within the SRM, and V657 and Y661 within the SET-I. D142 of the SRM electrostatically interacted with K660 of the SET-I, as described previously^18^. Further examination of the structural data revealed a potentially critical interaction between the SRM and EED during allosteric activation of PRC2. Moreover, several residues within the SRM, including H129 and H158, appeared to form an intercalated anchor at the EZH2-SRM/EED interface (Figure 1B, *right*). Intriguingly, while searching for genetic alterations of PRC2 subunits in human diseases using the COSMIC cancer mutation dataset and literature search, we found a number of mutations in the SRM/SET-I and SRM/EED interfaces that are present in human cancers and Weaver syndrome (Table S1). Of note, the EED mutations found in patients affected by Weaver Syndrome were exclusively located at the SRM/EED interface, further highlighting the potential role of this region in regulating PRC2 activity (Figure 1B, *right*, Table S1)^63–66^.

Taking into account the structural and human genetic data described above, we generated a series of point mutations within the SRM domain of EZH2, including P132S, D142V, and F145L which are located at the interface between the SRM/SET-I helices, as well as D136A that interacts with the Jarid2-K116me3 peptide (Figure 1B, *left*). Additionally, residues on EED that anchor the SRM/EED interaction were mutated and we termed them “EED anchoring mutants”, including H258Y, S259F, R302S, and R302G (Figure 1B, *right*). Versions of PRC2 carrying each of the aforementioned mutations were produced using a baculovirus expression system and purified through tandem affinity-tag purification (see Materials and Methods). The wild-type (WT) and mutant versions of PRC2 formed intact complexes (Figure S1) and importantly, exhibited similar levels of basal HMT activity (Figures 1C, 1E, 1G, and Figure S2). As expected, the EED^Y365A^ aromatic cage mutation specifically disrupted allosteric activation, and a catalytically inactive EZH2^(116-121A)^ mutant had neither basal nor stimulated activity (Figure 1C and Figure S3). Among the SRM and EED anchoring mutants described above, EZH2^P132S^, EZH2^F145L^ (within the SRM) and EED^R302G^ displayed a drastic reduction in allosteric activation, whereas EZH2^D136A^, EZH2^D142V^ (within the SRM), and other EED mutants rendered a moderate effect (Figure 1D, 1F, 1H and Figure S2). Structural modeling of the EED anchoring mutants indicated a disturbance in anchorage formation involving residues H129 and H158 of the SRM (Figure S4). Although the EED^Y365A^ aromatic cage mutant impaired allosteric activation of PRC2, the defect was due to its deficiency in interacting with H3K27me3, the initial recognition step, whereas other SRM and EED anchorage mutants did not manifest such a recognition defect (Figure S5), but were unable to be allosterically activated. Thus, two EZH2-SRM mutants (EZH2^P132S^ and EZH2^F145L^) and an EED anchoring mutant (EED^R302G^) are *bona fide* activation-deficient mutants of PRC2.

### Allosteric activation of PRC2/EZH2 is critical for appropriate H3K27me2/3 levels in vivo

Having defined the key residues of EZH2 and EED important for the response to H3K27me3-potentiated PRC2 allosteric activation *in vitro*, we next examined the role of these residues in regulating PRC2 activity *in vivo*. To this end, we generated EZH2^−/−^ (EZH2-KO) mESCs and transduced them with lentiviruses containing either EZH2^WT^, or the activation-deficient SRM mutants (EZH2^P132S^, EZH2^D142V^ or EZH2^F145L^). To study the EED anchoring mutations *in vivo*, we used a mESC line wherein the EED protein lacked the aromatic cage (see Materials and Methods). These cells (EED ∆C-term mESCs) were then transduced with lentiviruses containing EED^WT^ or EED anchoring mutants (EED^H258Y^, EED^S259F^, EED^R302S^, or EED^R302G^) (Figure 2A and 2B). While ectopic expression of EZH2^WT^ rescued H3K27me2 and H3K27me3 levels in the EZH2-KO mESC (Figure 2A, lane 4 and see Discussion), EZH2^P132S^ and EZH2^F145L^ were ineffectual (Figure 2A, lanes 5 and 7). On the other hand, EZH2^D142V^ displayed a partial rescue of H3K27me2, but the levels of H3K27me3 were severely compromised (Figure 2A, lane 6). In accordance, EED^WT^ and EED anchoring mutants regulated the endogenous levels of H3K27me2/3 levels in a manner similar to that observed *in vitro*, with EED^R302G^ (Figure 2B, lane 8) showing no rescue effect and EED^H258Y^, EED^S259F^, and EED^R302S^ showing a partial rescue of H3K27me2 levels, with levels of H3K27me3 being severely compromised (Figure 2B, lanes 5-7). These results were in full agreement with the biochemical studies for the stimulated activity shown above (Figures 1D, 1F, and 1H). Unexpectedly, these results revealed that the basal state of PRC2 had little impact on endogenous H3K27me2/3 levels and underscored the predominant role of allosteric activation of PRC2 in attaining appropriate levels of H3K27me2/3 *in vivo*. Indeed, mESC expressing EED^R302G^ (Figure 2B, lane 8) phenocopied those expressing EED ∆C-term and failed to form embryoid bodies (EB), a well-established phenotype associated with PRC2-deficiency^67–70^, as evidenced by the significantly low expression of the mesodermal marker, Brachyury (Figure S6). Of note, immunoprecipitation (IP) experiments using anti-EZH2 antibody demonstrated that the interactions between each of the SRM mutants and core PRC2 subunits as well as accessory proteins Jarid2 and Aebp2 were retained (Figure S7), excluding the possibility that these mutations destabilized PRC2 complex formation *in vivo*.

**Figure 2.**
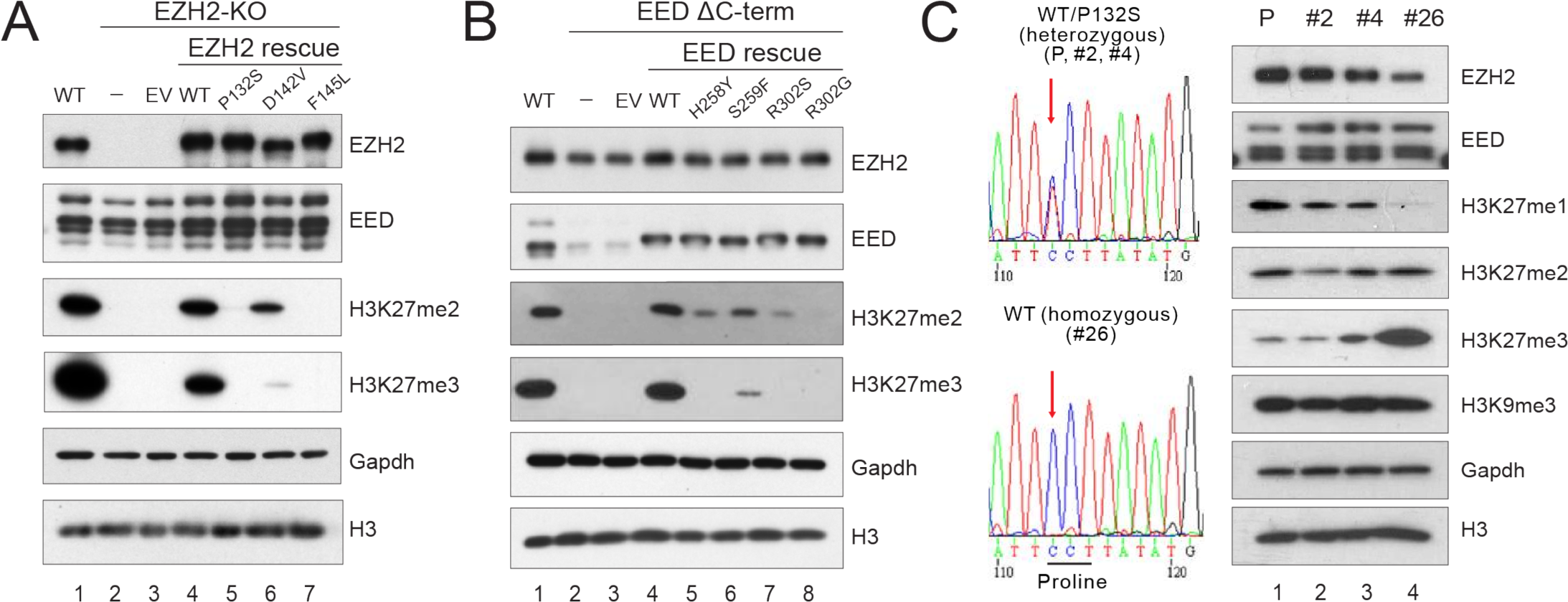
Point mutations of the key residues at the EZH2-SRM/SET-I or EZH2-SRM/EED interface affect global H3K27me2/3 levels *in vivo*. (A) Western blot analysis of EZH2, EED, H3K27me2, H3K27me3 and total histone H3 levels in E14 mESC cells, including WT, EZH2^−/−^ (EZH2-KO), and EZH2-KO cells rescued with EZH2^WT^ or EZH2-SRM mutants, as indicated. Gapdh was used as a loading control. mESC, mouse embryonic stem cells. EV, empty vector. (B) Western blot analysis of EZH2, EED, H3K27me2, H3K27me3, and total histone H3 levels in E14 mESC cells, including WT, EED∆C-term, and EED∆C-term cells rescued with EED^WT^ or EED anchoring mutants, as indicated. Gapdh was used as a loading control. EED∆C-term: EED C-terminal deletion. (C) CRISPR/Cas9-mediated reversal of EZH2^P132S^ to EZH2^WT^ in COLO-679 melanoma cells, which carry an endogenously heterozygous EZH2^P132S^ mutation. *Left*, Sanger sequencing results for the parental (P) COLO-679 cell line (EZH2^WT/P132S^), the negative CRISPR clones (EZH2^WT/P132S^; clones #2 and #4), and the positive CRISPR clone (EZH2^WT/WT^; clone #26). *Right*, Western blot analysis of EZH2, EED, H3K27me1, H3K27me2, H3K27me3, H3K9me3 and total histone H3 levels in the parental COLO-679 cells as well as CRISPR clones #2, #4, and #26. Gapdh was used as a loading control.

To further explore the impact of this positive feedback regulated allosteric activation of PRC2 in an independent system, we performed CRISPR/Cas9-mediated gene editing in a melanoma cell line, COLO-679, which carries a heterozygous EZH2^WT/P132S^ mutation. We generated a sub-clone (clone #26) that underwent a genetic “correction” and thus expressed only EZH2^WT^ from the parental line. This re-introduced EZH2^WT^ allele in clone #26 (EZH2^WT/WT^) led to markedly increased levels of H3K27me3, unchanged levels of H3K27me2, and decreased levels of H3K27me1 (Figure 2C, lane 4), when compared to the parental line (P, lane 1) and other negative CRISPR clones that remained EZH2^WT/P132S^ (clones #2 and #4, lanes 2 and 3, respectively). Of note, another repressive histone modification that is PRC2-independent, H3K9me3, was unaffected by the elimination of EZH2^P132S^ (Figure 2C). These results further corroborated the critical role of allosteric activation in driving the higher order methylation status of H3K27 by PRC2 *in vivo*.

### PRC2/EZH2 recruitment to chromatin is independent of its allosteric activation

As the allosteric activation-deficient mutants failed to rescue H3K27me2/3 levels *in vivo*, we next asked if these mutants impact PRC2 recruitment to chromatin. We performed chromatin immunoprecipitation followed by next-generation deep sequencing (ChIP-seq) using anti-EZH2 or anti-H3K27me3 antibody in WT and EZH2-KO mESCs, either alone or rescued with EZH2^WT^, EZH2^P132S^ or EZH2^F145L^. Ectopic expression of EZH2^WT^ in the EZH2-null background rescued the majority of the EZH2 peaks across the genome and at the transcription start sites (TSS) of PRC2/EZH2 target genes (Figure 3A and 3B, *left*). Surprisingly, the allosteric activation deficient mutants, EZH2^P132S^ and EZH2^F145L^, not only rescued the genome-wide occupancy, but also manifested a more pervasive association with chromatin in either the max-peak-centered or TSS-centered heatmap (Figure 3A and 3B, *left,* see Discussion). Yet, while each mutant bound to chromatin, H3K27me3 levels remained similar to the case of EZH2-KO cells (Figure 3A and 3B, *right*). Of note, we observed an incomplete rescue of the genome-wide H3K27me3 peaks upon ectopic expression of EZH2^WT^ in EZH2-KO cells, possibly due to clonal variation, as our EZH2-KO line was a clonal line derived from the CRISPR/Cas9 approach (Figure 3A, *right*). However, in the same experiment, EZH2^WT^ did rescue most of the H3K27me3 peaks at the TSS of PRC2/EZH2 targets, suggesting that H3K27me3 occupancy at gene promoters is more conserved and fluctuated less between the clonal and the parental lines (Figure 3B, *right*). Intriguingly, we detected a small number of H3K27me3 peaks in EZH2-KO cells (Figure 3A-C), likely due to the presence of EZH1 that functions similarly, but to a considerably lesser extent than EZH2^16,23, and Lee et al, in submission^. Regardless of the residual H3K27me3 peaks detected at a small number of loci (Figure 3C), EZH2^WT^, EZH2^P132S^, and EZH2^F145L^ were still fully deposited to the same loci as endogenous EZH2 in control E14 mESC, suggesting that the chromatin landscape in EZH2-KO mESCs preserved an epigenetic memory that facilitated appropriate restoration of PRC2/EZH2 occupancy (Figure 3A-C). These results indicated that PRC2 recruitment to target genes in the chromatin context is uncoupled from its enzymatic activity.

**Figure 3.**
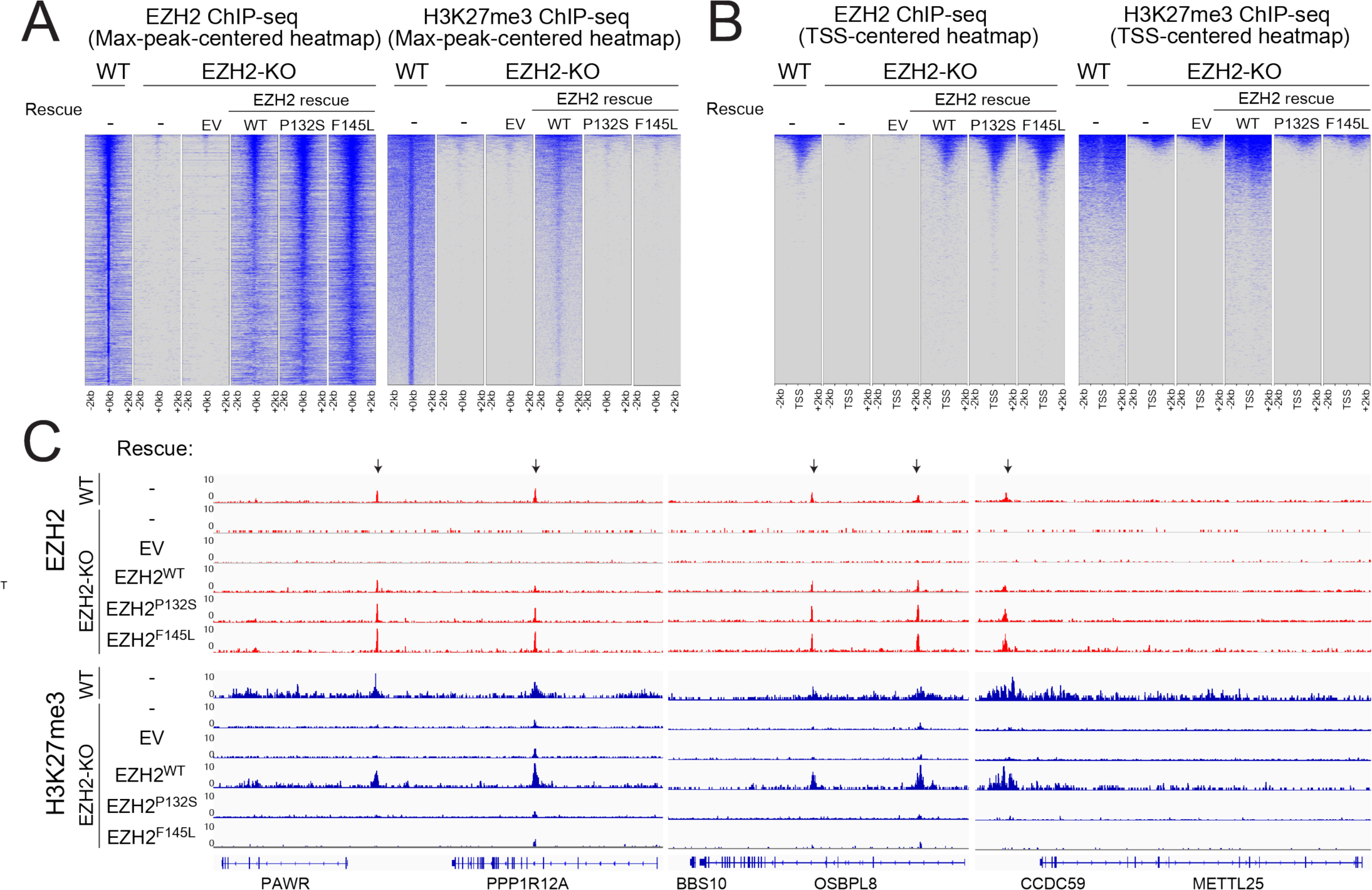
The activity of PRC2 is independent of its recruitment to chromatin. (A and B) Heatmaps representing EZH2 (left panels) or H3K27me3 (right panels) ChIP-seq peaks centered by maximum peak intensity in the genome-wide scale (A) or ~29000 UCSC annotated transcription start sites (TSS) and (B) in 4-kb windows within 200 bp bin size. The genotypes of E14 mESC for each experimental condition are indicated above each panel. WT, wild-type. KO, knockout. EV, empty vector. (C) Representative track images for EZH2 (in red) and H3K27me3 (in blue) ChIP-seq experiments performed in panel (A and B) are shown for a select group of annotated genes. The UCSC annotations of exons and gene bodies are shown at the bottom. Arrows indicate the MACS peaks with FDR <5% in the respective windows. The scale of the peaks ranges from 0-10 reads per 10 million reads, with spike-in normalization using *Drosophila* chromatin and *Drosophila* H2A.X antibody in each ChIP reaction.

### An allosteric activation-deficient mutation epistatically overrides a gain-of-function mutation in EZH2

As allosteric activation appears to be the predominant mechanism governing PRC2/EZH2 catalysis *in vivo*, we speculated that this mechanism might override the effect of cancer-associated, gain-of-function mutations at the Y646 residue within the SET domain of EZH2. Therefore, we generated a compound mutant of EZH2 that carried both the P132S activation deficient mutation (in the SRM) and the Y646N gain-of-function mutation (in the SET domain). We measured the HMT activity of PRC2 comprising EZH2^WT^, EZH2^P132S^, EZH2^Y646N^, or EZH2^P132S;Y646N^ using mono-, di-, and un-methylated H3K27 oligonucleosomes as templates, as a function of the presence of the allosteric activator (H3K27me3). Consistent with the previous peptide-based HMT assay, PRC2 comprising EZH2^Y646N^ exhibited no basal activity on un-methylated or mono-methylated H3K27 templates (Figure 4A, lane 4 and Figure 4B, lane 7). Moreover, EZH2^Y646N^ had higher basal activity compared to EZH2^WT^ on di-methylated templates (Figure 4C, compare lanes 1 and 7). This is consistent with previous findings that the gain of function mutation (EZH2^Y646X^) exhibited an altered substrate preference for H3K27me2 over H3K27me1 or unmethylated templates^47,49^. In accordance, EZH2^Y646X^ oncogenic mutations were always found to be heterozygous, requiring one EZH2^WT^ allele to cooperatively acquire a gain-of-function state by providing the H3K27me2 substrate^71^. More importantly, EZH2^Y646N^ was also regulated by allosteric activation as it catalyzed elevated levels of H3K27me3 in response to the H3K27me3 peptide (Figure 4C, lanes 7-9). However, the presence of this allosteric activator was ineffectual in the case of EZH2^P132S;Y646N^ (Figure 4C, lanes 10-12), suggesting that the gain-of-function effect driven by EZH2^Y646N^ was restrained at the basal level by disruption of the induced interaction between SRM/SET-I.

**Figure 4.**
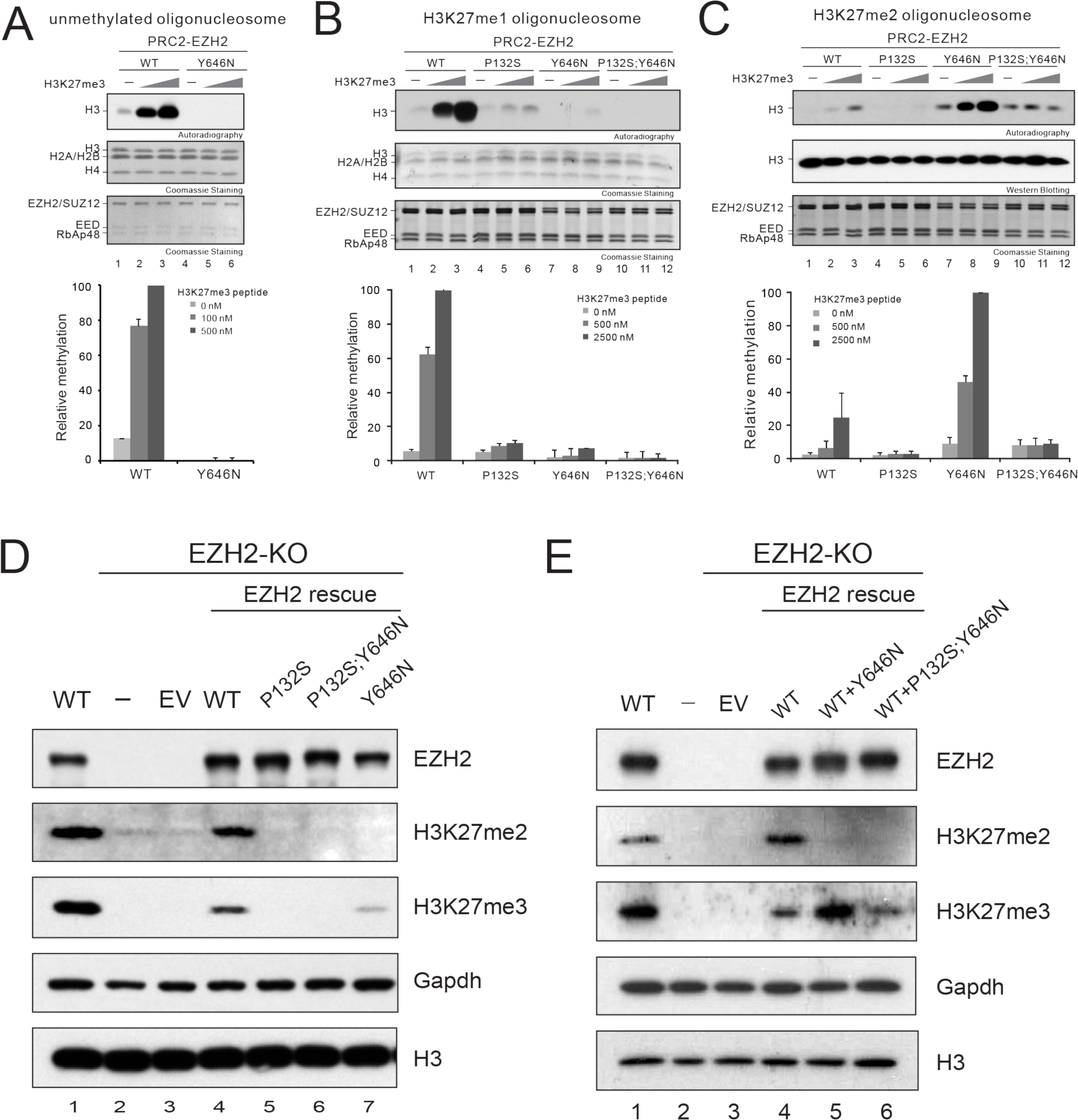
EZH2^P132S^-driven inhibition of the allosteric activation of PRC2 overrides the effect of the EZH2^Y646N^ gain-of-function mutation. (A-C) Histone methyltransferase assays. The details of the HMT assay conditions as well as the generation of the oligonucleosomes are described in Materials & Methods. (A) *Top*, a representative image of the HMT assays containing 100 nM WT PRC2 or PRC2 containing EZH2^Y646N^ with unmodified oligonucleosomes (150 nM) as substrate in the absence or presence of H3K27me3 peptide (100 or 500 nM). The levels of methylation on histone H3 are shown by autoradiography. Coomassie blue staining of SDS-PAGE gels containing nucleosomes (middle image) or PRC2 components (bottom image) was used to visualize the relative concentration of each component present in each reaction. *Bottom*, quantification of the relative amounts of ^3^H-SAM incorporated into histone H3 (n=3 for each data point). (B-C) *Top*, representative images for the HMT assays containing 500 nM WT PRC2 or PRC2 containing EZH2^P132S^, EZH2^Y646N^, or EZH2^P132S;Y646N^ in the absence or presence of the H3K27me3 peptide (500 or 2500 nM) using 150 nM of H3K27me2 oligonucleosomes (B) or 150 nM of H3K27me1 oligonucleosomes (C) as substrate. The concentrations of PRC2 and H3K27me3 peptide were adjusted in these HMT reactions due to the poor catalytic activity of PRC2-EZH2^Y646N^ on H3K27me1 or H3K27me2 oligonucleosomes. The middle/bottom gel images and the bottom panel for loading controls and signal quantifications, respectively, are as described in (A). (D) Western blot analysis of EZH2, H3K27me2, H3K27me3, and total histone H3 in WT, EZH2-KO, and EZH2-KO E14 mESCs rescued with EZH2^WT^, EZH2^P132S^, EZH2^Y646N^, or EZH2^P132S;Y646N^. Gapdh was used as a loading control. EV, empty vector. (E) Western blot analysis of EZH2, H3K27me2, H3K27me3, and total histone H3 levels in WT, EZH2-KO, and EZH2-KO E14 mESCs rescued with EZH2^WT^ alone or co-rescued with EZH2^Y646N^ or EZH2^P132S;Y646N^. Gapdh was used as a loading control. EV, empty vector.

To test if PRC2/EZH2^Y646N^ catalysis of H3K27me3 *in vivo* was dependent on its allosteric activation, EZH2^Y646N^ and EZH2^P132S;Y646N^ were stably expressed in EZH2-KO mESC using a lentivirus approach. As expected, expression of EZH2^Y646N^ alone in EZH2-null cells only partially rescued H3K27me3 due to a lack of EZH2^WT^ that compromised the catalysis of mono- and di-methylation of H3K27 (Figure 4D, lane 7). Nevertheless, similar to EZH2^P132S^, EZH2^P132S;Y646N^ was ineffectual in rescuing H3K27me3 levels, demonstrating that the activity of the hyperactive mutant was dependent on allosteric activation *in vivo*, and regulated in a manner similar to that of WT PRC2 (Figure 4D, lanes 5 and 6). To recapitulate the heterozygous nature of the EZH2^Y646N^ mutation found in cancers, we co-expressed EZH2^WT^ with either EZH2^Y646N^ or EZH2^P132S;Y646N^ in EZH2-null cells. In the presence of EZH2^WT^, EZH2^Y646N^ manifested its expected hyperactivity for the catalysis of H3K27me3 (Figure 4E, lane 5), whereas EZH2^P132S;Y646N^ was devoid of such hyperactivity (Figure 4E, lane 6). Thus, the P132S mutation epistatically overrode the effect of the Y646N gain-of-function mutation in catalyzing H3K27me3 *in vivo*.

### Disrupting allosteric activation of PRC2/EZH2 effectively inhibits proliferation of PRC2/EZH2-addicted tumors

Next, we sought to evaluate the impact of allosteric activation of PRC2 on PRC2/EZH2-dependent human cancers. We focused on DLBCL cell lines that harbor EZH2^Y646X^ mutations, as the majority of EZH2^Y646X^-mutant cell lines showed a dependency on the activity of EZH2 and several SAM-competitive inhibitors are in clinical trials in Follicular Lymphoma (FL) and DLBCL. We compared two EZH2^Y646X^-mutant DLBCL cell lines: KARPAS-422 (EZH2^WT/Y646N^), a known PRC2/EZH2-dependent cell line and SU-DHL-4 (EZH2^WT/Y646S^), an exceptional case without such EZH2-dependency. By using a lentivirus-based system, we first stably introduced an shRNA against the 3’ untranslated region (3’-UTR) of endogenous EZH2 mRNA transcripts in both cell lines. As expected, knockdown of endogenous EZH2 showed a drastic reduction in H3K27me3 levels in both cell lines (Figure 5A, lanes 2 and 6), but only inhibited proliferation of KARPAS-422 cells (Figure 5B, compare black and blue lines). Next, we performed rescue experiments in the EZH2-knockdown background of each cell line. Ectopic co-expression of EZH2^WT^ and EZH2^Y646N^ by lentiviruses restored H3K27me3 levels (Figure 5A, lane 7) and proliferation of KARPAS-422 cells (Figure 5B, *right*, red line), demonstrating that the effects of the EZH2 shRNA was on-target. However, rescue by co-expression of EZH2^P132S^ and EZH2^P132S;Y646N^ failed to negate the effects of EZH2 knockdown in these cells (Figure 5A, lanes 4 and 8, and Figure 5B). On the other hand, co-expression of EZH2^WT^ and EZH2^Y646N^ partially rescued H3K27me3 levels in the SU-DHL-4 cells, likely due to the insufficient restoration of EZH2 levels (Figure 5A, lane 3). Nevertheless, concomitant expression of EZH2^P132S^ and EZH2^P132S;Y646N^ was ineffectual with respect to H3K27me3 levels and neither of the rescue conditions had any effect on SU-DHL-4 cell proliferation, due to their PRC2/EZH2-independency (Figure 5A and 5B, *left*). These proof-of-principle experiments highlighted the requirement of PRC2 allosteric activation for H3K27me3 catalysis and the dependency on the oncogenic activity of Y646-mutant EZH2 for proliferation of those DLBCL cells that are EZH2-dependent. These findings point to the importance of the SRM/SET-I interface as a potential target in a therapeutic strategy against cancers that rely on either WT or hyperactive PRC2/EZH2.

**Figure 5.**
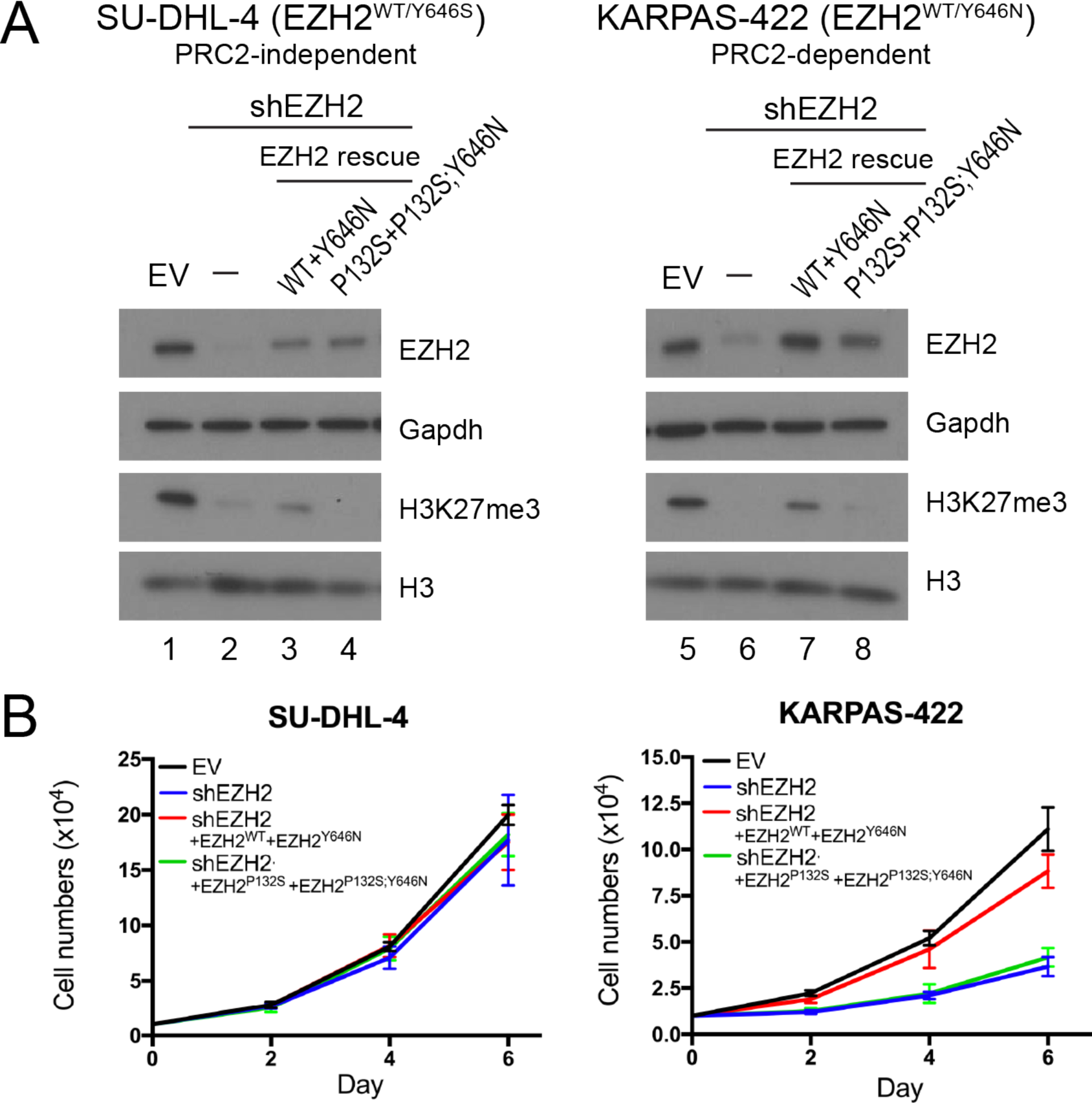
Genetic inactivation of the allosteric activation of PRC2 impairs both H3K27me3 levels in EZH2^Y646X^-mutant DLBCLs and proliferation of PRC2-addicted cells. (A) Western blot analysis of EZH2, H3K27me3, and total histone H3 levels in SU-DHL-4 (left, PRC2-independent) and KARPAS-422 (right, PRC2-independent) DLBCL cells with control (EV), EZH2 knockdown (shEZH2), and EZH2 rescue conditions, as indicated on the right side. Gapdh was used as a loading control. EV, empty vector. (B) Proliferation assays for SU-DHL-4 (left) and KARPAS-422 (right) cells under the abovementioned conditions. 10,000 cells were seeded in 6-well plates and cell numbers were counted after 2, 4, and 6 days. The culture medium was replenished every two days. Cell numbers are shown as mean ± SD (n=3 for each data point).

### Disrupting EZH2-SRM/SET-I interaction by alpha-helical mimetics selectively inhibits allosteric activation of PRC2

As shown above, the interaction between the SRM and SET-I domain of EZH2 is mediated by two closely aligned α-helical structures. To disrupt this protein-protein interaction, we utilized a well-established oligopyridylamide scaffold-based α-helix mimetic approach. The oligopyridylamides form a network of intramolecular hydrogen bonds that stabilize a single conformation and project side chain functionalities from one face that correspond to the side chain residues of an α-helical conformation at i, i+3 (i+4), and i+7^72^. The use of these oligopyridylamides has been shown to effectively disrupt α-helix-mediated protein-protein interactions^73,74^. To disrupt the SRM/SET-I helical interaction, we synthesized a library of oligopyridylamides by designing the surface functionalities complementary to the side chain residues of the SRM helical interface (Figure 6A and 6B). For example, to mimic side chain residues of the SRM helix (residues D142, F145, and L149), we designed oligopyridylamides ADH-61 (side chain groups: Amine, Benzyl, Benzyl). We conducted a structural activity relationship study to further our understanding of the mode of action of the oligopyridylamides against the SRM/SET-I interface and ADH-61 was predicted to be one of the most potent antagonists. We postulated that the amine group on ADH-61 forms a salt bridge with the carboxylate group of D142 and that the two benzyl groups on ADH-61 interact with the hydrophobic side chains (F145 and L149) of the SRM helix (Figure 6A and 6B, *left*). ADH-24 served as a negative control in which the amine group was replaced with carboxylate and the central planar benzyl group was replaced with a non-planar cyclohexane in order to diminish its hydrophobicity as well as the salt bridge interaction with the SRM helix (Figure 6A and 6B, *right*). We applied these α-helix mimetics to the HMT assays, in the absence or presence of the allosteric activator (H3K27me3 peptide). As predicted, incubation with ADH-61 selectively inhibited reconstituted WT PRC2 from responding to the stimulatory H3K27me3 peptide, whereas ADH-24 had no overt effect (Figure 6C). The half maximal inhibitory concentration (IC_50_) of ADH-61 was 0.766 ± 0.710 μM for inhibition of PRC2 HMT activity *in vitro* (Figure 6D). Moreover and importantly, neither ADH-61 nor ADH-24 exhibited an overt effect on the basal activity of PRC2 (Figure 6E). The data indicate that ADH-61 likely inhibited PRC2 HMT activity by targeting the SRM/SET-I interface. We also investigated the effect of ADH-61 on the toxic-gain-of-function of PRC2/EZH2^Y646N^ mutant. Using H3K27me2 oligonucleosomes as template, the gain-of-function of PRC2/EZH2^Y646N^ towards H3K27 tri-methylation was efficiently inhibited by ADH-61 in a dose-dependent manner with an IC_50_ of 0.891 ± 0.096 μM, while ADH-24 had no significant effect on the PRC2/EZH2^Y646N^ mutant (Figures 6F and 6G). Together, these results provided the proof-of-principle for the concept that ordering of the SRM and the ensuing SRM/SET-I interaction are indeed central to PRC2 allosteric activation.

**Figure 6.**
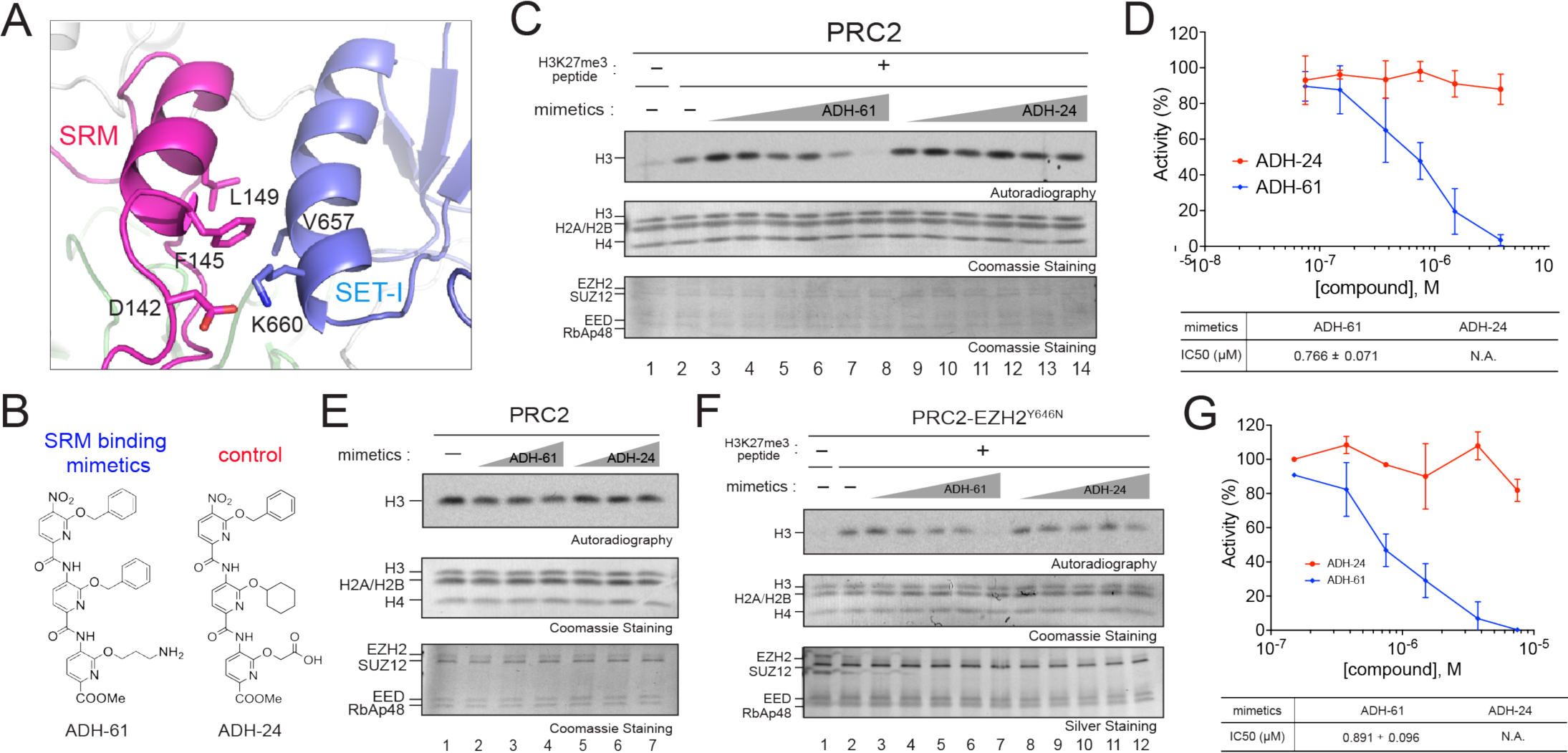
Oligopyridylamide-mediated selective inhibition of the allosteric activation of PRC2. (A) An enhanced image depicting molecular features of the EZH2-SRM/SET-I interface. Key residues on the interface of SRM and SET-I α-helices are highlighted. (modified from PDB:5HYN) (B) Chemical structures of the α-helix mimetics used in the study. ADH-61 was designed to interact with the amino acid side chains containing D142, F145, and L149 of the EZH2-SRM domain facing the SET-I domain. (C) Representative HMT assays containing 7.5 nM PRC2 in the absence or presence of the H3K27me3 peptide (150 nM), with increasing amounts (0.075, 0.15, 0.375, 0.75, 1.5 or 3.75 µM) of the α-helix mimetic compounds using unmodified oligonucleosomes (150 nM) as substrate. Details of the HMT assay conditions are described in Materials & Methods. Autoradiography shows the levels of methylation on histone H3 (top) and Coomassie blue staining of SDS-PAGE gels shows histones (middle) or PRC2 components (bottom). (D) *Top*, Quantification of the relative amounts of ^3^H-SAM incorporated in the HMT assays described in (C) (n=3 for each data point). *Bottom*, IC_50_ was extracted for ADH-61 using a dose dependent study. (E) Representative images of the HMT assays containing 7.5 nM PRC2 with increasing amounts (0.75, 1.5 or 3.75 µM) of the α-helix mimetic compounds in the absence of the H3K27me3 peptide using unmodified oligonucleosomes (150 nM) as substrate. Other panels are as described in (C). (F) *Top*, Representative images of HMT assays containing 10 nM PRC2 comprising of EZH2^Y646N^ with increasing amounts (0.15, 0.375, 0.75, 1.5, or 3.75 µM) of ADH-61 or ADH-24 in the absence or presence of the H3K27me3 peptide (300 nM) using H3K27me2 oligonucleosomes (150 nM) as substrate. Other panels are as described in (C). (G) Quantification of the relative amounts of ^3^H-SAM incorporated in the HMT assays described in (F) (n=3 for each data point). *Bottom*, IC_50_ was extracted for ADH-61 using a dose dependent study.

## Discussion

Mammalian heterochromatin contains large repressive chromatin domains, including H3K9me2/3-decorated constitutive heterchromatin at pericentromeric and telomeric regions and H3K27me2/3-decorated facultative heterochromatin with a more context-dependent deposition^75,76^. The writers of these two classes of histone modifications include Suv39h1/2 and PRC2^34,77,78^, respectively. These enzymes are regulated by a “read-and-write” mechanism whereby they recognize their own catalytic product through a chromodomain (CD) and an aromatic cage of EED, respectively. Such recognition, in turn, stimulates their enzymatic activity as a positive feedback loop that is determinant for the spreading and the formation of the respective repressive chromatin domain^34,77–79^.

Although the exact mechanism of allosteric activation of the enzymes that catalyze H3K9 methylation are not clearly defined, the studies presented here provide a detailed mechanism of H3K27me2/3-mediated allosteric activation by PRC2. The EED core subunit of PRC2 binds to H3K27me2/3 *in vitro* and *in vivo*^34^, and initiates a dynamic reorganization of the structure of the core PRC2 complex comprising EED, SUZ12 and EZH2. Such reorganization following H3K27me2/3 recognition by the EED-cage results in a positive feedback stimulatory loop exemplified by the promotion of a stable interaction between the SRM and SET-I α-helices of EZH2 as well as an interaction between the SRM and EED anchorage region. This allosteric stimulation-induced reorganization not only promotes catalysis, but also sustains a read-and-write mechanism facilitating the spreading and formation of H3K27me2/3-decorated repressed chromatin domains.

By identifying PRC2 mutants that specifically abrogate its allosteric activation, we demonstrated that this effect is crucial for its activity *in vivo* and the requirement for its basal activity observed *in vitro* may not be met *in vivo*, at least in mESCs (Figures 1 and 2). Our findings also highlighted an unprecedented, unique role of EZH2-SRM/EED anchorage in addition to the EZH2-SRM/SET-I interface in mediating the allosteric effect on H3K27 methylation. This notion is further supported by the EED anchorage mutations found in patients affected by Weaver Syndrome (Table S1 and Figure S4)^63–66^. Unlike the initial discovery of EED cage mutants, the EZH2-SRM and EED anchoring mutants are not defective in H3K27me3 binding (Figure S5), allowing us to specifically segregate the allosteric effect and dissect the stepwise action of PRC2. An interesting observation from our study is that the EZH2-SRM and EED anchoring mutants manifested a spectrum of inhibitory effects on the allosteric activation of PRC2, with several of them retaining the ability to catalyze H3K27me2 *in vivo* (Figure 2A and 2B). These results implicate the depletion of H3K27me3, but not that of H3K27me2, as being key to the development of those diseases associated with these respective mutations. Further analysis of mutants specifically defective in H3K27me3 catalysis should be performed to elucidate the potentially unique role of H3K27me3 in these contexts.

Inherently, EZH2 catalyzes H3K27me2 more efficiently than H3K27me3 (Figure 4A-4C), as the Y646 residue within the SET domain likely restricts the transition from H3K27me2 to H3K27me3^80^. Intriguingly, the gain of function EZH2^Y646X^ oncogenic mutations were shown to break this rate-limiting step by switching substrate preference and therefore, specifically promoting the conversion of H3K27me2 to H3K27me3. Such a deregulation of the catalytic kinetics is central to the oncogenesis in DLBCL and melanoma, underscoring the importance of the rate-limiting control of the feed-forward mechanism and balancing endogenous H3K27me3 levels at undesired loci underlying disease pathogenesis^47–49,80,81^. Importantly, our results demonstrated that the allosteric effect not only governs the activity of WT PRC2 *in vivo*, but also surpassed the hyperactivity of PRC2 containing the EZH2^Y646N^ mutation (Figures 4D and 4E). In PRC2/EZH2-addicted DLBCL cells, genetic inactivation of the allosteric effect suppressed their proliferation as well as their endogenous H3K27me3 levels (Figure 5). Together with the proof-of-principle experiments exploiting the α-helix mimetics against the EZH2-SRM/SET-I interaction (Figure 6), these results provided a molecular and functional basis for future development of a different class of allosteric inhibitor in addition to the recently developed allosteric inhibitor that binds to the aromatic cage of EED^57,58,60^. Albeit challenging, a number of protein-protein interaction (PPI) inhibitors have been developed and are under clinical testing, such as the BCL2-BH3 domain inhibitor, ABT-199 and the MDM2/TP53 interaction inhibitor, RG7112^82–86^. In addition, the interfaces of EZH2-SRM/SET-I and EZH2-SRM/EED contained a groove surface feature (Figure S8), which may offer ideal target sites for the screening or structural design of potential small molecular inhibitors against the allosteric activation of PRC2.

To date, a number of factors have been implicated in regulating PRC2 recruitment to chromatin, perhaps reflective of the distinct types of genes targeted by PRC2 and that PRC2 is, by default, a general repressor of transcription, ensuring the maintenance of a repressed state^20,87^. At imprinted genes or at active protein coding genes, the main PRC2 recruiter appears to be non-coding RNA (ncRNA) or the nascent transcript itself, respectively^88–91^. We suggest that at these genes, PRC2-mediated repression occurs passively, as silencing of gene expression is intially established by other factors (such as DNA metylation or sequence-specific repressors, respectively) prior to PRC2 recruitment to target DNA sequences. Therefore, deposition of H3K27me2/3 at these regions is a slow process and likely less dependent on H3K27me2/3-mediated allosteric activation. However, at other genes, such as those that regulate lineage commitment, the role of PRC2 is much more active. Upon initial PRC2 recruitment, the deposition of H3K27me2/3 is critical and highly dependent on its allosteric activation. Recruitment can be mediated by accessory factors known to associate with PRC2, such as Jarid2, Aebp2, and PCLs^20,23,29,32,33,88^, as well as DNA itself ^(Lee et al, submitted)^. Among these factors, Jarid2 perhaps plays a pivotal role. Jarid2 interacts with PRC2 independently of chromatin, but their binding to chromatin at CpG islands is inter-dependent. Given that PRC2 methylates Jarid2 at lysine 116 (Jarid2K116me3), which functions similarly to H3K27me3, we speculate that Jarid2K116me3 is important in igniting the allosteric mechanism upon PRC2 recruitment to its target genes for *de novo* H3K27me3 deposition.

In addition, a very interesting and surprising finding from our study is the uncoupling of PRC2 recruitment and catalysis in the context of chromatin (Figure 3). We observed that the EZH2-SRM mutants displayed increased and more pervasive association with chromatin as compared to WT EZH2, implying that the activated state of PRC2 might restrict its local occupancy independently of its initial recruitment. Is the actual deposition of H3K27me2/3 important in restricting/delineating PRC2-mediated repressed domains? If this were to be the case, as the data suggests, then H3K27me2/3 has yet an unknown function that perhaps requires the action of other factors.

In summary, our data defined the intermediate state during the allosteric activation of PRC2, which involved a structural remodeling and interplay between EZH2 and EED. We also demonstrated that the allosteric effect of PRC2 is a dominant feature in catalyzing H3K27 methylation in the context of chromatin comprising genes at which PRC2 is a major determinant in establishing an effective repressed state. Yet surprisingly, its allosteric activation can be uncoupled from PRC2 recruitment. As drug resistance against SAM competitive EZH2 inhibitors remains a future challenge, a new class of allosteric inhibitors targeting the EED-cage has recently been developed. Our findings on the suppression of EZH2^Y646X^ hyperactivity and DLBCL proliferation by abrogating allosteric activation underscores the possibility of devising a novel class of allosteric inhibitors that target the SRM/SET-I interaction or SRM/EED anchorage against PRC2/EZH2-addicted cancers.

---

**Figure S1. PRC2 containing EZH2-SRM or EED anchoring mutants form intact complexes.** Purified recombinant PRC2 (EZH2, EED, SUZ12 and RbAp48) containing WT EZH2, EZH2-SRM activation-deficient mutants, EZH2 gain-of-function mutants, or EED anchoring mutants. Proteins were detected by Coomassie blue staining of SDS-PAGE gels.

**Figure S2. PRC2 containing EZH2-SRM or EED anchoring mutants showed impaired responsiveness to the H3K27me3 peptide for the stimulated activity but no defect for the basal activity.**

(A-C) Histone methyltransferase assay results with loading controls performed as in Figure 1C-1H. The levels of methylation on histone H3-containing oligonucleosomes are shown by autoradiography (Top image) and Coomassie blue staining of SDS-PAGE gels containing nucleosomes (middle image) or PRC2 components (bottom image) was used to visualize the relative concentration of each component present in each reaction.

**Figure S3. SAL (SET-activation loop) of EZH2.** SAL (yellow) interacts with EED (green), SET domain of EZH2 (blue), and VEFS domain of SUZ12 (brown). The side chain of amino acids 116-121 in SAL is shown and PRC2 containing six alanine mutations on these residues, EZH2^(116-121)A^, is catalytically inactive (see Figure 1). (modified from PDB:5HYN)

**Figure S4. Structural modeling of EED anchorage mutants found in Weaver Syndrome.**

(A) Structural depiction of the EZH2-SRM/EED interface. Critical residues that mediate interactions were highlighted and the distance among residues were measured using Pymol program and indicated.

(B-D) Structural modeling of EED^R302G^ (panel B), EED^R302S^ (panel C), or EED^H258Y^ (panel D) was shown. EED^R302G^ and EED^R302S^ mutation lose its interactions with EZH2^H129^ and EZH2^H158^. On the other hand, EED^H258Y^ mutation causes steric collision with the main chain of EZH2^H158^ (See the distance between two residues). (modified from PDB:5HYN)

**Figure S5. PRC2 containing EZH2^P132S^ or EZH2^D136A^ binds to H3K27me3 peptide.** Immunoprecipitation (IP) using a H3K27me3 peptide (amino acids 18-36) for the pull-down of WT PRC2 or PRC2 containing EED^Y365A^, EZH2^P132S^, or EZH2^D136A^ mutants. Coomassie blue staining of input and IP samples separated on a SDS-PAGE gel was shown to visualize protein abundance. The EED cage mutant (EED^Y365A^) manifested a binding defect to the peptide, whereas EZH2-SRM mutants (EZH2^P132S^ and EZH2^D136A^) showed no defect.

**Figure S6. mESCs expressing PRC2/EED^R302G^ phenocopies the PRC2-deficiency phenotype of embryoid body (EB) differentiation.**

Western blot analysis of Brachyury, H3K27me3 and total histone H3 levels in WT or EED∆C-term mESCs rescued with EED^WT^ or EED^R302G^ after 3 days of EB differentiation, as well as undifferentiated WT mESCs. The whole cell extract of mESCs was prepared as described in Materials and Methods. Brachyury was used as a mesodermal marker. EV, empty vector.

**Figure S7. EZH2-SRM mutants form intact complexes together with other PRC2 core subunits and accessory proteins *in vivo*.** Immunoprecipitation (IP) using an anti-EZH2 monoclonal antibody (D2C9 Cell signaling) in WT and EZH2-KO E14 mESCs rescued with the indicated constructs. The mESC nuclear extract was prepared as described in Materials and Methods. Input and IP samples were separated on an SDS-PAGE gel. Western blot analysis for the PRC2 core subunits (EED and SUZ12) and co-factors (Jarid2 and Aebp2) is shown. Asterisk indicates a background band for the heavy chain of the antibody used in EZH2-IP. Total histone H3 was used as a loading control for input samples.

**Figure S8. The EZH2-SRM/SET-I interface and the EZH2-SRM/EED anchorage contain groove structures.**

(A-B) Surface representation of EZH2-SRM (pink)/SET-I (blue) interface. The positions of D142, F145, and P132 of EZH2 are highlighted in red. The groove pockets are indicated by black arrows. (C) Structural animation of EED. The positions of R236, H258, and R302 of EED are highlighted in dark grey. The groove feature generated by R236, H258, and R302 of EED is indicated by a black arrow. (D) Surface representation of EZH2-SRM (red)/EED (grey) anchorage. H129 and H158 of EZH2 are highlighted in pink, and R236 and R302 of EED are highlighted in dark grey. (modified from PDB:5HYN)

## Materials and Methods

### DNA constructs

To purify human PRC2 core complexes, FLAG-tagged-EED, 6xHIS-tagged-EZH2, SUZ12, and RbAp48 were cloned into a baculovirus expression vector, pFASTBac1 (Invitrogen). EZH2 or EED mutant constructs were generated by site-directed mutagenesis and mutations were confirmed by Sanger DNA sequencing. The EZH2^(116-121A)^ mutant consists of six alanine mutations on the SAL (SET-activation Loop) domain of EZH2, thereby preventing the establishment of a stable conformation of the SET domain^17^. WT or mutant EZH2 and EED constructs were subcloned into the pLV-EF1-alpha-IRES-mCherry vector (Clontech) for lentiviral production and delivery.

### Antibodies

Antibodies against EED and Jarid2 were produced in-house. Other commercial antibodies against EZH2 (Cell signaling, D2C9, catalog no. 5246), SUZ12 (Cell signaling, D39F6, catalog No. 3737), Aebp2 (Cell signaling, D7C6X, catalog No. 14129), H3K9me3 (Millipore, catalog no. 07-442), H3K27me1 (Millipore, catalog no. 07-448), H3K27me2 (Cell signaling, D18C8, catalog no. 9728), H3K27me3 (Cell signaling, C36B11, catalog no. 9733), H3 (Abcam, catalog no. ab1791), Gapdh (GeneTex, catalog no. GTX627408), were purchased from the companies indicated.

### Purification of protein using Baculovirus expression system

All four core components of PRC2 (FLAG-EED, 6xHIS-EZH2, SUZ12, and RbAp48) were co-expressed in Sf9 cells by baculovirus infection. After 60 hr of infection, Sf9 cells were resuspended in BC150 buffer (25 mM Hepes-NaOH, pH 7.8, 1 mM EDTA, 150 mM NaCl, 10 % glycerol, 1 mM DTT and 0.1 % NP-40) with protease inhibitors (1 mM phenylmethlysulfonyl fluoride (PMSF), 0.1 mM benzamidine, 1.25 mg/ml leupeptin and 0.625 mg/ml pepstatin A) and phosphatase inhibitors (20 mM NaF and 1 mM Na_3_VO_4_). Cells were lysed by sonication (Fisher Sonic Dismembrator model 100), and WT or mutant recombinant PRC2 was tandemly purified through Ni-NTA agarose beads (Qiagen), FLAG-M2 agarose beads (Sigma), Q sepharose beads (GE healthcare) and glycerol gradient sedimentation (15-35% glycerol gradient).

### Nucleosome reconstitution

Recombinant histones were generated as previously described^16^. Briefly, each core histone was expressed in BL21 (DE3) Codon Plus cells (Agilent), extracted from inclusion bodies, and purified by sequential anion and cation chromatography. For refolding recombinant octamers, equal amounts of histones were mixed and dialyzed into refolding buffer (10 mM Tris-HCl, pH 7.5, 2 M NaCl, 1 mM EDTA, and 5 mM β-mercaptoethanol). Octamers were further purified by size exclusion chromatography on a 24-mL Superdex 200 column (Amersham Biosciences) in refolding buffer. Recombinant oligonucleosomes were reconstituted by sequential salt dialysis of octamers and plasmid having 12 601-nucleosome positioning sequences. The methyl-lysine analogue production for the generation of pseudo-monomethyl- and pseudo-dimethyl-H3K27 nucleosomes was previously described in detail^11^.

### HMT assay

Standard HMT assays were performed in a total volume of 15 μL containing HMT buffer (50 mM Tris-HCl, pH 8.5, 5 mM MgCl_2_, and 4 mM DTT) with 500 nM of ^3^H-labeled S-Adenosylmethionine (SAM, Perkin Elmer), 10 nM (500 ng) of recombinant oligonucleosomes consisting of 12x repeat nucleosome arrays (120 nM of nucleosome), and recombinant human PRC2 complexes. The reaction mixture was incubated for 60 min at 30 °C and stopped by the addition of 4 μL SDS buffer (0.2 M Tris-HCl, pH 6.8, 20% glycerol, 10% SDS, 10 mM β-mercaptoethanol, and 0.05% Bromophenol blue). A titration of PRC2 (from 5 to 60 nM) was performed under these conditions in order to optimize the HMT reaction within a linear range, and the yield of each HMT reaction was measured using the following procedures. After HMT reactions, samples were incubated for 5 min at 95 °C and separated on SDS-PAGE gels. The gels were then subjected to Coomassie blue staining for protein visualization or wet transfer of proteins to 0.45 μm PVDF membranes (Millipore). The radioactive signals were detected by exposure on autoradiography films (Denville Scientific). To measure “basal” HMT activity of PRC2, a titration of WT PRC2 or PRC2 harboring EZH2 or EED mutations was used at a concentration of 7.5, 15, or 30 nM. To measure the “stimulated” HMT activity of PRC2, 15 nM of PRC2 or PRC2 harboring EZH2 or EED mutations was used in the presence of an H3K27me3 peptide (amino acids 21-44) at a concentration of 0, 50, or 250 nM. All HMT assays were performed with the above indicated concentrations of PRC2 and H3K27me3 peptide unless otherwise noted in the figure legends.

### Cell culture and EB formation

Mouse embryonic stem cells (mESCs) were grown in standard ESC medium containing Lif, 1 μM MEK1/2 inhibitor (PD0325901) and 3 μM GSK3 inhibitor (CHIR99021). For EB formation, 600K mESCs were plated in suspension plates with medium containing DMEM, 20% FBS, 1% Pen/Strep, 2 mM L-Glutamine, and 0.1 mM 2-mercaptoethanol, 1 mM vitamin C. Cells were collected 3 days after the induction of EB differentiation.

### CRISPR/Cas9-mediated genome editing

An optimal gRNA target sequence closest to the genomic target site was chosen using the http://crispr.mit.edu/ design tool. The gRNA was cloned into the SpCas9-2A GFP (Addgene: PX458) or SpCas9-2A Puro (Addgene:PX459) via BbsI digestion and insertion^92^. mESC cells were seeded into 12-well plates at 100,000 cells per well, and transfected with 0.5 μg of the appropriate guide RNAs, template DNA for guide RNAs (listed in Table 2) and Cas9 endonuclease using Lipofectamine 2000 (Life Technologies). The transfection was performed according to the manufacturer’s recommended protocol, using a 3:1 ratio of Lipofectamine:DNA. After transfection, GFP-positive cells were sorted using the Sony SY3200 cell sorter, and 20,000 cells were plated on a 15 cm dish. Puromycin resistant cells were selected in 2 μg/mL puromycin for 48 hr 7-10 days later, single ESC clones were picked, trypsinized in 0.25 % Trypsin-EDTA for 5 min, and plated into individual wells of a 96-well plate for genotyping. Genomic DNA was harvested via QuickExtract (Epicentre) DNA extraction, and genotyping PCRs were performed using primers surrounding the target site. The resulting PCR products were purified and sequenced to determine the presence of a deletion or a mutation event. EED C-term deletion mESCs were generated by a frame-shift deletion resulting in a stop codon at residue D371 of EED.

### Lentiviral production and delivery

For the production of viral particles, lentiviral vectors containing WT or mutant EZH2 and EED (10 μg) were co-transfected with pcREV (2.5 μg), BH-10 (3 μg), and pVSV-G (5 μg) packaging vectors into 293-FT cells. The virus-containing medium was collected 48 hr after transfection and the target cells were spin-infected. Polybrene was added to the viral medium at a concentration of 8 μg/mL. Infected cells were FACS-sorted for mCherry.

### Synthesis of α-helix mimetics

The monomers for the preparation of the oligopyridylamides were synthesized according to a previously published method^73^. Following the monomer synthesis, chain elongation of pyridylamides was achieved using iterative amide coupling between oligopyridylamines and monomeric-pyridylacids using 2-chloro-1-methylpyridinium iodide (Mukaiyama’s reagent) followed by reduction of the nitro groups. The acid labile tert-butyl esters and NH-*Boc* groups were cleaved using a trifluoroacetic acid (TFA) cocktail (dichloromethane/TFA/triethylsilane, 80:15:5, v/v/v) in the final step to attain the series of oligopyridylamides.

### ChIP-seq and data analysis

Standard ChIP experiments were performed as previously described^93^. 100 µg sonicated chromatin was used in each ChIP reaction with 2.5 µg of anti-EZH2 or anti-H3K27me3 antibody. 1 µg of *Drosophila* chromatin and 0.1 µg of anti-*Drosophila* H2A.X antibody were added in each ChIP reaction as spike-in references. For ChIP-seq library preparation, up to 30 ng of immunoprecipitated DNA was end-repaired, A-tailed, and ligated to custom barcode adapters with T4 ligase as previously described^93^. Libraries were sequenced on Illumina HiSeq. A custom barcoding system was employed. The ChIP-seq data were first mapped to the mouse genome (mm9 downloaded from UCSC genome browser) using the Bowtie2 software package (version 2.3.0). All reads that failed to align to the mouse genome were mapped to the fly genome (dm6 downloaded from UCSC genome browser) using the same software and criteria. The sum of the number of reads mapped to both genomes was used as the actual “library size”. The median of the library sizes across all the samples was used as the reference library size, which is about 20,000,000 reads. All samples were scaled to obtain the new library size equal to the reference library size. The input samples were normalized to the same total reference library size as the ChIP samples. After adjustment of the total library size, the number of reads mapped to the fly genome was used as the spike-in normalization factor. All samples were then scaled except for input samples in order to have the same total spikein content. To obtain EZH2 binding sites, only reads with mapping quality greater than 30 were selected and removed as presumptive PCR duplicates with the same mapped coordinates. MACS2 software package was used (version 2.1.1) for calling significantly enriched peaks at a false discovery rate (FDR) less than 5% relative to the input samples. For H3K27me3 samples, SICER software package was used (version 1.1) to find broad peaks by setting the window size of 500 bp and allowing peaks within a distance of 500 bp to merge together. The significantly enriched peaks were filtered with the same criteria as above, FDR < 0.05.

### Preparation of mESC whole cell extract

8×10^6^ cells were harvested and lysed with standard RIPA buffer (10 mM Tris-HCl (pH 7.5), 150 mM NaCl, 1mM EDTA, 1% Triton X-100, 0.1% sodium deoxycholate, and 0.1% SDS) containing protease inhibitors (0.2 mM PMSF, 1 µg/mL Pepstatin A, 1 µg/mL, Leupeptin, and 1 µg/mL Aprotinin) and phosphatase inhibitors (10 mM NaF and 1 mM Na_3_VO_4_). The cell suspension was briefly sonicated (40% amplitude, 5 strokes) and centrifuged at 20,000 x g at 4 ºC for 20 min. The supernatant was collected as the whole cell extract.

### Preparation of mESC nuclear extract

To prepare nuclear extract from mESC, approximately 4×10^7^ cells were harvested and washed with PBS. Cells were lysed with intact nuclei in TMSD buffer (40 mM Tris-HCl, pH 7.5, 5 mM MgCl_2_, 250 mM sucrose, 1 mM DTT, and 0.02% NP-40) containing protease inhibitors and phosphatase inhibitors. The cell suspension was centrifuged at 800 x g at 4 °C for 10 min. The pellets (nuclei) were resuspended in 20x cell pellet volume of BC400 buffer (20 mM Tris-HCl, pH 7.9, 400 mM KCl, 0.2 mM EDTA, 20% glycerol, 0.5 mM DTT, and 0.02% NP-40) with protease inhibitors and phosphatase inhibitors, and incubated at 4 °C for 1 hr. The cell suspension was centrifuged at 18,000 x g at 4 ºC for 20 min. The supernatant was collected as the nuclear extract.

### Immunoprecipitation

2 mg of mESC nuclear extract was incubated with 3 µg of corresponding antibody at 4 ºC for 2 hr. 0.6 mg of Dynabeads Protein G (Thermo fisher) was added and incubated at 4 ºC for 1 hr. The tubes were placed on magnets to separate the beads from the solution and unbound supernatant was removed. The beads were washed three times with BC400 buffer at 4 ºC for 10 min. After the final wash, samples were incubated for 5 min at 95 °C to extract protein from the beads and separated on SDS-PAGE gels. The gels were then subjected to wet transfer of proteins to 0.45 μm PVDF membranes for Western Blotting.

### Peptide binding assay

2 nmol of biotinylated H3K27(18-36) or H3K27me3(18-36) was immobilized to Streptavidin agarose resin (Millipore). 2 µg of recombinant PRC2 or PRC2 containing EED or EZH2 mutants was added and incubated at 4 ºC for 1 hr. Unbound supernatants were removed and beads were washed three times with BC150 at 4 ºC for 10 min. After the final wash, samples were incubated for 5 min at 95 °C to extract protein from the beads and the recovered protein was separated on SDS-PAGE gels. The gels were then subjected to Coomassie blue staining.

## Author contributions

C-H.L., J-R.Y., and D.R. conceptualized and designed the study. C-H.L, J-R.Y., and S. Kaneko conducted the experiments. S.K. designed and synthesized the α-helical mimetics in A.D.H’s lab. Y.J. performed bioinformatics analyses. J-R.Y, C-H.L, and S.K. wrote the manuscript with assistance from D.R.

## Acknowledgement

We thank Drs. L. Vales, R. Margueron, and K-J. Armache for critical reading of the manuscript as well as past and current Reinberg members for critical comments and discussion; D. Hernandez, O. Oksuz, K. Stafford, and J. Stafford for technical assistance. We also thank the NYU Flow Cytometry Core (grant: NIH/NCI P30CA016087) for cell purification. The work in D.R.’s lab is supported by NIH (NCI R01CA199652) and the Howard Hughes Medical Institute (HHMI). The work in A.D.H’s lab is supported by New York University. J-R.Y. is supported by the American Cancer Society (PF-17-035-01).

